# A Tale of Two Environments: Divisive Normalization and the (In)Flexibility of Choice

**DOI:** 10.1101/2024.08.25.609561

**Authors:** Vered Kurtz-David, Shreya Sinha, Vinayak Alladi, Stefan Bucher, Adam Brandenburger, Kenway Louie, Paul Glimcher, Agnieszka Tymula

## Abstract

The Divisive Normalization (DN) function has been described as a “canonical neural computation” in the brain that achieves efficient representations of sensory and choice stimuli. Recent theoretical work indicates that it efficiently encodes a specific class of Pareto-distributed stimuli. Does the brain shift to different encoding functions in other types of environments, or is there evidence for DN encoding in other types of environments? In this paper, using a within-subject choice experiment, we show evidence of the latter. Our subjects made decisions in two distinct choice environments with choice sets either drawn from a Pareto distribution or from a uniform distribution. Our results indicate that subjects’ choices are better described by a divisive coding strategy in both environments. Moreover, subjects appeared to calibrate a DN function to match, as closely as possible, the actual statistical properties of each environment. These results suggest that the nervous system may be constrained to use divisive representations under all conditions.

**Significance Statement:** How does the frequency with which we encounter different kinds of decision problems affect how the brain represents those problems? Recent empirical findings suggest that we adapt our internal representations to match the environments in which we are making choices. Theoretical work has shown that one form of internal representation, called divisive normalization, provides an optimal adaptation when making choices in a specific class of environments. Using a stylized experimental design, subjects faced two distinct choice environments, each characterized by different statistical properties. Our findings show humans appear to use the same mechanism in both environments, suggesting that a divisive representation may be a fixed feature of human cognition.

## Introduction

We make some decisions more often than others – in dozens of instances during our life we choose between having a pizza or a burger for dinner, but rarely have to indicate which of the two-starred Michelin restaurants we prefer. An often overlooked fact is that these encounter frequencies play a critical role in defining efficient encoding strategies – given constraints on neural coding, more accurate encoding must generally be allocated to more frequently encountered stimuli (1, 2). Indeed, experimental studies confirm this theoretical insight, showing a dependency of preference orderings, choice patterns (3–7), and choice efficiency (3, 8) on the frequency with which subjects encounter different rewards.

This has led to the conclusion that during the decision process, the brain adheres to principles of *efficient coding*, economizing the allocation of resources to optimize decision outcomes (3, 8–13). A canonical example of a well-studied efficient code (13, 14) is Divisive Normalization (henceforth DN) (15), which has been related to neuronal firing rates across all sensory modalities (16–19) and across various cognitive domains as well (20). The DN function enables a system with limited information capacity to employ a flexible and scale-invariant encoding of naturally occurring stimuli that is sensitive to encounter frequency (17, 21, 22). Ample evidence has supported the notion that DN is also highly predictive of reward value encoding in the human and animal choice mechanism (6, 7, 23–25).

At least one form of DN has been shown to be an efficient code for stimuli with a probability of occurrence that is described by a Pareto Type III distribution (26). This prompts the question of whether the brain employs non-DN encoding functions when the statistical properties of the input stimuli (in our case, *choice environment*s) are not Pareto-distributed. Would we expect to find evidence of divisive encoding mechanisms (13) – like cross-normalized DN (26) – only in Pareto-distributed environments? The latter might imply that previous documentation of DN encoding mechanisms may say more about the antecedent stimulus distributions used in experiments than about constraints on encoding mechanisms. An alternative hypothesis, however, is that our brains are constrained to employ DN-like encoding mechanisms (13). Such a constraint might reflect an adaptation of the nervous system to Pareto-distributed real-world natural stimuli, such as the sensory (14, 18, 27) and ecological (28, 29) environments we typically encounter.

In this study, we present our subjects with a binary-choice task in two environments characterized by different reward distributions. In one environment, valuations are Pareto-III-distributed and hence at least some DN functions are efficient (13, 26). In the other, valuations are uniformly distributed. In such an environment, a DN encoder would not represent the environment most efficiently (26) (Figure 1A). We test hypotheses about our subjects’ value encoding functions by fitting the patterns of errors in their choices with two random utility models (henceforth, RUM) (30, 31). The first one is a function that captures the key features of the family of DN models (32, 33). The second is a RUM with a power utility function that is the standard model in economic research and lies outside the DN family^1^ (henceforth, power utility; Figure 1B). We test our data to determine which model, divisive or non-divisive, better describes subjects’ choices in each environment. We use a generalized form of DN to examine whether subjects are better described as obligate-DN choosers who use DN in both environments, or alternatively, that subjects’ choices are better be described with our DN function in one environment and with a power utility function in the other (Figure 1C).

**Figure 1.**
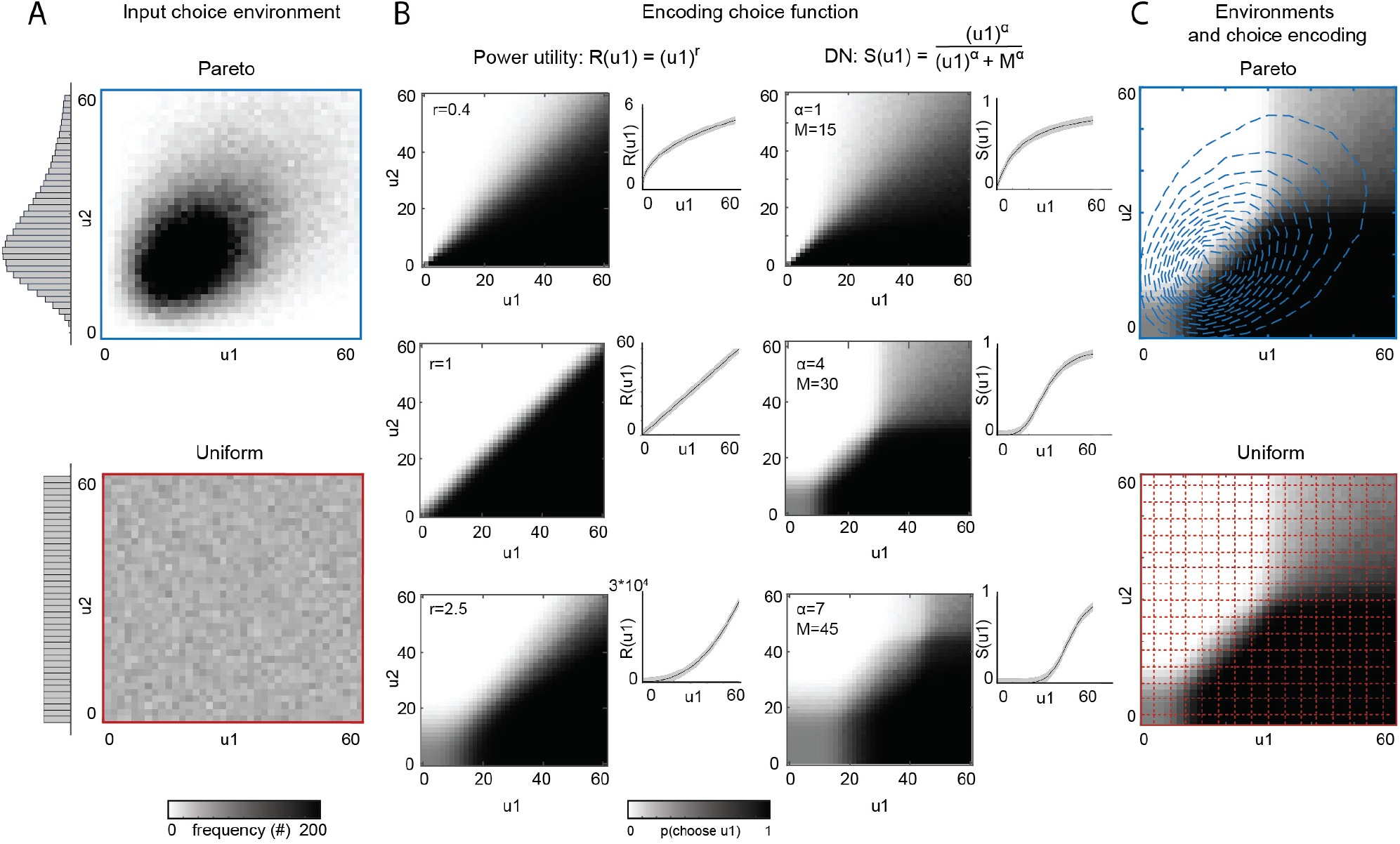
Research Question. (A) Choice environments are determined by the distribution of valuations. We compare a long-tailed bivariate Pareto Type III environment for which DN is efficient code with a uniformly distributed environment for which DN is not an efficient code. Figures show 2D histograms of simulated choice trials with valuations in the range *u*_*k*_ ∈ [0,60] for every lottery *k* ∈ {1,2}. Each reward’s value was drawn from 40 bins. Insets show their corresponding marginal distributions. We simulate 100k valuations per environment. See Materials & Methods for further details. (B) Value encoding choice functions. We test two different RUM models: classic power utility (left) and DN (right). The figure shows the probability of choosing a lottery with valuation *u*_1_ over a lottery with valuation *u*_2_ for various parameter values in each model. Insets show the subjective representation of *u*_1_ in power utility (*R*), and in DN (*S*). For every combination of *u*_1_ and *u*_2_, we simulate 1k binary choice sets. We allow stochasticity in choice by incorporating additive noise, drawn from 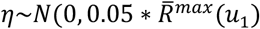, such that 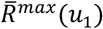 denotes the maximal subjective value of *u*_1_ in the power utility model (and 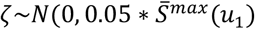 in the DN model, respectively). We cast 10K noisy draws per simulated trial and reported average choice probabilities across simulated sets. (C) Contour plots indicate the mass of occurrences of (*u*_1_, *u*_2_) choice trial combinations in each environment. Contours were laid over a representative DN model with *α* = 4, *M* = 30 (middle right panel in (B).

In line with theory (13, 26), we find that in a Pareto-distributed environment, subjects employ an encoding of values that is well modeled by DN. We also find that the DN model better captures subjects’ choices in the uniformly-distributed environment. This suggests that subjects’ choices are more accurately described by divisive encoders, like those found in DN models, than by standard power utility functions. We find further evidence for context dependency in subjects’ choices as within the constraints of DN encoding, they adapt their reward expectations according to changes in the specific statistical properties of the choice environment.

Taken together, our results suggest that divisive mechanisms may be an obligate component of the encoding mechanism used during the choice process. The current study focuses on decision making processes, but given the dominance of DN representations across cortical systems, our findings may be of general interest to the study of encoding mechanisms in sensory and other cognitive domains.

## Results

### Two-Stage Task Design

Seventy-six subjects completed a two-stage choice task. In STAGE I (Fig. 2A left panel), subjects reported their valuations (willingness to pay) for 33 50-50 lotteries that pay either *y*_1_ or *y*_2_ dollars (see Table S1 for a complete lottery list). These valuations were used to estimate, for every subject *i*, their STAGE I subjective value function 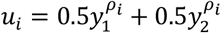, using a standard non-linear least squares (NLS) estimation. The subjective value function curvature (*ρ*_*i*_) varied substantially from subject to subject (Fig. 2E). Using individual *ρ*_*i*_ estimates, we generated subject-specific distributions of rewards in terms of their subjective – rather than dollar – values for the STAGE II task (Fig. 2B). This first step was critical. It allowed us to perform all our analyses in the domain of subjective value, removing simple utility curvature from our primary analyses and allowing us to create individualized choice sets with specific distributional properties that were essential for our design. Without this transformation, small subject-specific differences in utility curvature (risk attitudes) would have made the construction of probative choice sets required for the experiment impossible.

**Figure 2.**
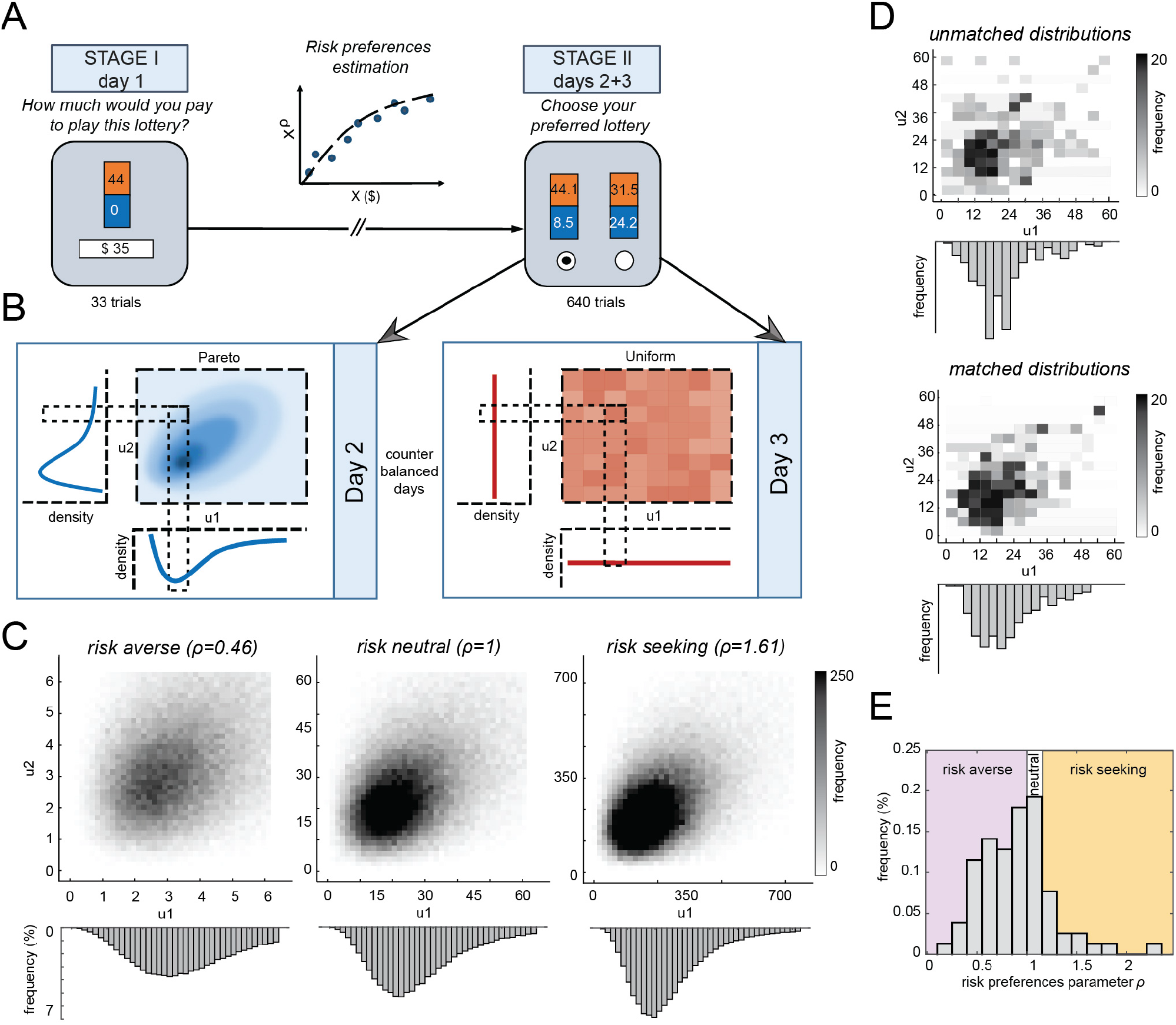
Experimental Design. (A) Timeline. In STAGE I, subjects reported their valuations for 33 lotteries. Valuations were used to recover the curvature of the subjective value function for each subject using NLS estimation. Based on those estimates, we generated subject-specific bi-dimensional uniform and Pareto Type III distributions of valuations for STAGE II of the study. In STAGE II, subjects completed two sets of 320 binary choices between 50-50 lotteries (640 choices in total). (B) Bi-dimensional Pareto and uniform distributions. In the uniform distribution, we created 40 bins of subjective values between 0 and the maximal payoff in the study 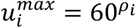 with eight lotteries in each bin. We then picked pairs of lotteries from this set to create binary choice sets. In the Pareto distribution, we used a Gamma-weighted scale mixture of exponential random variables to capture the covariance structure of the bi-variate Pareto distribution. (C) Choice sets in STAGE II controlled for differences in individual subjective value function (risk attitudes), modulating the second moment (std) of the Pareto distribution (see eq. (vi) in Materials and Methods). The histograms show the bi-dimensional Pareto distributions and their marginals (with 100k draws per distribution) from three representative subjects: left - a risk averse subject, middle - a risk neutral subject, right - a risk seeking subject. (D) Experimental sets with 320 trials were prone to under-sampling (see top, unmatched distribution). We matched experimental sets to the distributional shape of a larger set with 100k draws (see bottom, matched distributions). The figure shows an example corresponding to the middle panel in (C). (E) Recovered estimates of subjective value curvature (risk attitudes) from STAGE I. See Methods and Figure S1 for further details. See Table S2 for a list of the estimated subjective value function curvatures (risk parameter *ρ*).

In STAGE II, on two separate days, subjects made binary choices between 50-50 lotteries (Fig. 2A, right panel), with 320 decisions on each day. We created two choice environments: on one day, subjects were choosing between lotteries with subjective values drawn from a Pareto Type III distribution, for which DN has been proven to be an efficient encoder (henceforth, Pareto), and on the other day between lotteries with subjective values drawn from a uniform distribution for which DN has been shown to not be an efficient encoder (26). Subjects encountered each distributional environment on a different day (counter-balanced across subjects) to avoid contextual spillovers.

Using these risky-choice lotteries, rather than choices over consumer goods, enabled us to generate *continuous* distributions of valuations for STAGE II and to fully control their *distributional shape*. As noted above, our decision to generate the distributions of STAGE II lotteries in subjective value space, rather than in dollar space, aimed to control for the heterogeneity in subjects’ subjective valuations of lotteries (risk attitudes, Fig. 2E and Table S2). If, instead, we had created these distributions based on the dollar amounts, then for most subjects (all those with a non-linear subjective value function, i.e. *ρ*_*i*_ ≠ 1) STAGE II choice sets would have not corresponded to uniform or Pareto type III distributions due to the heterogeneity in their subjective valuation of monetary lotteries. Our two-stage procedure thus ensured that for all subjects the environments had the same distributional shape – enabling us to assess the effect of the distributional properties of the two choice environments, while fully controlling for individual differences in preferences.

Across subjects, we fixed the first moment (mean) of valuations and the range of monetary payoffs in both environments. Naturally, the second moment (standard deviation) of the uniform distribution was also fixed across subjects. The second moment of the Pareto distribution (as measured in dollars) varied by subjects’ subjective valuations of money as assessed in STAGE I (risk attitudes) (Fig. 2C, Fig. S1).

Accordingly, this heterogeneity also varied the distributions of the high and low monetary payoffs in each lottery (Fig. S1). To ensure that we fully captured each distributional environment, we matched the mean and standard deviation of the choice sets with those of larger sets of 100k draws (Fig. 2D). See Materials and Methods for further details on our sampling design.

Overall, subjects appeared to pay careful attention during the study – only six subjects in the uniform environment, and nineteen subjects in the Pareto environment failed to choose the higher subjective value lottery in more than 20% of trials (Fig. S2A). Respectively, on average, subjects violated first-order stochastic dominance in 0.97% of trials in the uniform treatment and in 1.08% of trials in the Pareto treatment (Fig. S2B). Note that a higher incidence of mistakes in the Pareto environment is expected – as due to the correlational structure across lotteries, the value difference between lotteries was (on average) smaller and thus choices were harder in the Pareto environment (34).

### Distributional Properties of the Choice Environments Influence Subjects’ Choice Behavior

Our overarching goal was to study how the distributional properties of the choice environment influenced the encoding of value, and whether subjects could flexibly switch between different types of encoding mechanisms, as evidenced by errors in their choice patterns, in different environments. We created the experimental choice environments with Pareto Type III and uniform distributions of valuations. In this section, we tackle the first part of our research question in a model-free manner, determining whether the distributional structure influenced the errors produced by our subjects in a meaningful manner.

It is useful to introduce our hypotheses using an illustration. In Fig. 3A-B, we indicate the probability of choosing lottery 1 with valuation *u*_1_, given the coupling of the (*u*_1_, *u*_2_) valuations in a choice set. Choices along the diagonal represent trials in which the two lotteries had the same or very similar valuations, whereas trials that are away from the diagonal correspond to choice sets in which the two lotteries’ valuations were substantially different. A central feature of DN is the calibration of the function to the input stimuli. That is, resources are allocated to the range of stimuli most likely to be observed (*tuning*) (15, 25). Thus, compared to non-divisive encoders, if DN governs the choice mechanism in a Pareto environment, we would expect that in this environment subjects would make more mistakes in choice sets with elements away from the high-density center of the main diagonal, because these choices are less frequent. Conversely, we also expect that subjects in the Pareto environment would make fewer mistakes in choices whose valuations lie near the main diagonal because these near-equivalued choices occur more frequently. We find both patterns in our data.

**Figure 3.**
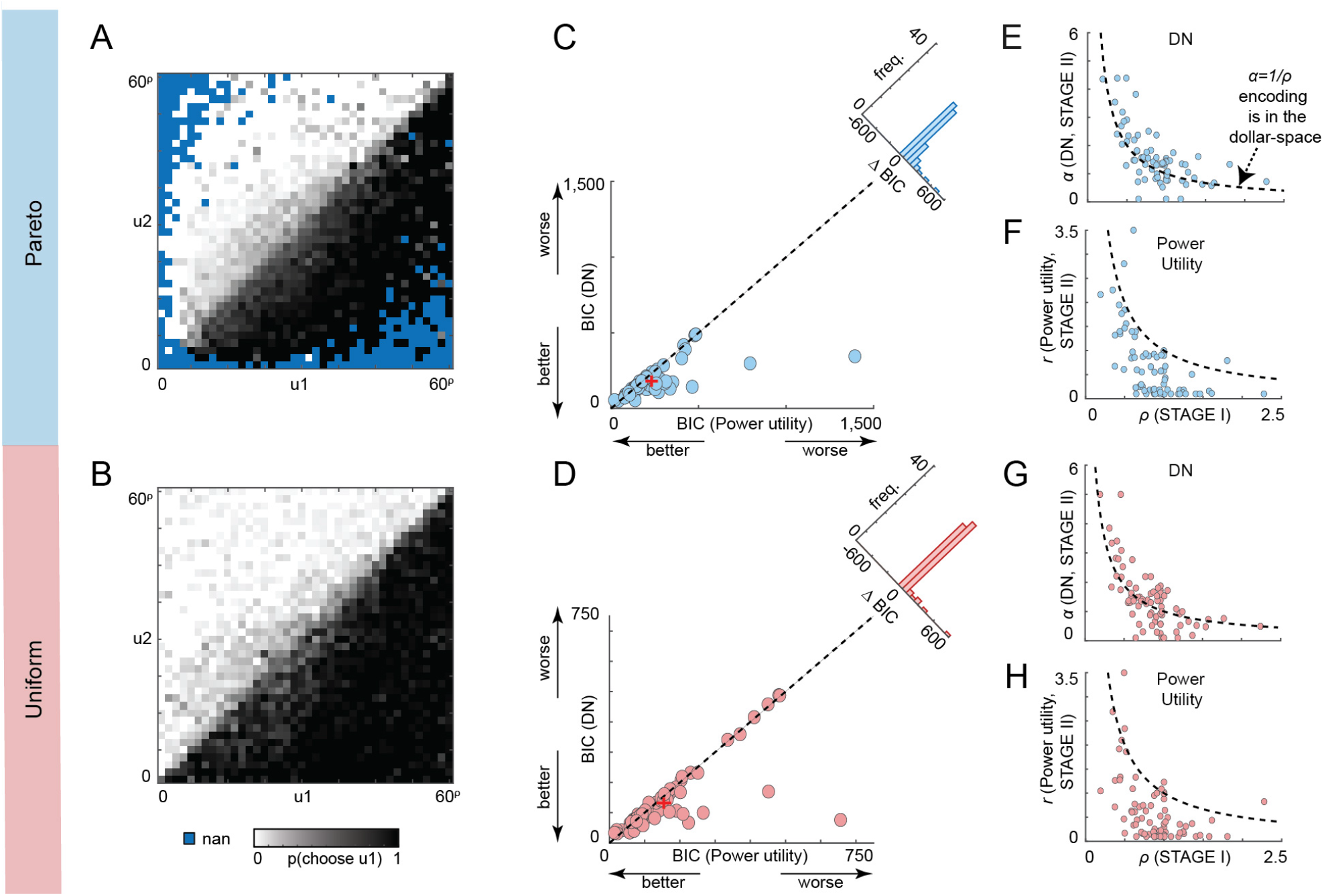
Model-fitting. (A-B) Probability of choosing lottery 1 with valuation *u*_1_, in a (*u*_1_, *u*_2_) choice set. Data is aggregated over subjects. Within subjects, valuations are divided into 60 equally-spaced bins. (A) The Pareto environment. (B) The uniform environment. (C-D) Each dot is one subject’s DN model BIC score (y-axis) plotted against the same subject’s power utility BIC score (x-axis). A dashed 45-degree line indicates when both models are equally successful. Inset shows the difference in BIC scores (*BIC*_*power*_ - *BIC*_*DN*_. (C) The Pareto environment. (D) The uniform environment. (E-F) Relationship between the STAGE I curvature of the subjective value function (ρ) and STAGE II subjective value functions in the Pareto environment. Dashed curve indicates hyperbolic function 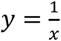. (E) DN model (*α* parameter). (F) Power utility model (r parameter). (G-H). Same as (E-F), but for the uniform environment. Dots indicate individual subjects, + indicate the sample averages. N=76.

To statistically test whether the frequency of mistakes increased faster as choice sets moved away from the main diagonal in the Pareto environment than in the uniform environment, we ran a probit regression with an indicator dependent variable equal to one for trials on which a subject selected the option with higher SV, and equal to zero otherwise. We controlled for the difference in difficulty across the trials by including the absolute value difference between the lottery valuations (|*u*_1_ − *u*_2_|) and for the general impact of the distribution by including a dummy for the Pareto distribution. The different rate of mistakes depending on the distance from the diagonal in each environment are captured by a significant coefficient on the interaction of Pareto dummy and (|*u*_1_ − *u*_2_|) (Column (1) in Table 1). As expected, we found that choice accuracies increased with an increase in the subjective value distance between the two options, and that moving from the uniform distribution to the Pareto distribution reduced accuracy (see also discussion in the previous section). Importantly, in line with our hypothesis, we found a negative and significant interaction term, indicating that relative to the uniform distribution, in the Pareto distribution, subjects were more likely to make errors once encountering choice sets further away from the diagonal, those sets that they experienced less often in the Pareto environment. We can thus conclude that encounter frequency as defined by the Pareto distributional structure did influence choice accuracy.

**Table 1.** Results from the model-free analysis. Probit regressions with the dependent variable is equal to 1 when the subject chose the lottery with the higher SV, and zero otherwise. Column (1) model was run on the full sample. The independent variables are the absolute SV difference between the two lotteries, a dummy indicating the Pareto environment and their interaction. Column (2) model was run on data including choice sets in the center of the distributions. The model includes the same independent variables as model (1), and an additional dummy equal to 1 if the lottery was taken from around the diagonal (and zero otherwise, see text for definitions) and its interaction with the Pareto dummy. Standard errors clustered on subject in parentheses, + p<0.1, * p<0.05, ** p<0.01, *** p<0.001.

|  | (1)<br>Full<br>sample | (2)<br>Center of the<br>distributions |
| --- | --- | --- |
| SV difference | 0.0002*<br>(0.0001) | -0.0001<br>(0.0000) |
| Pareto | -0.3311***<br>(0.0405) | -0.1479***<br>(0.0374) |
| Pareto*SV difference | -0.0001*<br>(0.0001) | -0.0003***<br>(0.0001) |
| Near diagonal |  | -0.7400***<br>(0.0692) |
| Pareto*Near diagonal |  | 0.1742**<br>(0.0597) |
| Constant | 1.3260***<br>(0.0704) | 1.1201***<br>(0.0713) |
| N | 48640 | 22442 |
| pseudo R-sq | 0.015 | 0.036 |

To examine whether subjects calibrated their encoding function to the most frequently presented choice sets, we tested if they made fewer mistakes around the high-density center of the main diagonal in the Pareto environment. We ran a complementary probit regression focusing on twenty-two valuation bins from the center of the distributions presented in Fig. 3A-B (out of an equally-spaced 40-bin space), which corresponded to lotteries with $9-42 payoffs^2^ (Column (2) in Table 1). In addition to the regressors used in the model above, we included a dummy variable that indicated whether a lottery was taken from around the diagonal of the valuation space, as well as its interaction with the Pareto distribution dummy. We defined lotteries as laying around the diagonal if the ratio between the two valuations was 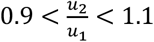.

Not surprisingly, choice accuracy was lower in choice sets around the diagonal, since these represented the most difficult choices in the experiment with the smallest SV difference. Crucially though, we found a positive interaction term between the diagonal and Pareto dummies, suggesting that relative to the uniform environment, in the Pareto environment, subjects had higher accuracy in those particularly difficult trials within the highly sampled region. Our results remained robust for other definitions of *the center of the distribution* and *around the diagonal* (Table S3).

Together, these results suggest that in the Pareto environment subjects adjusted their value encoding to increase choice accuracy rates at the center of the joint distribution, at the expense of the decreased choice accuracy at the margins. This is evidence for a divisive form of value encoding, where choice discriminability is the highest near the mode of the distribution (see Fig. 1B).

### Evidence for DN-like Value Encoding Across Choice Environments

The findings in the previous section provided initial evidence that subjects adapted to the distribution of valuations and that subjects used divisive encoding in the Pareto environment. Our next goal was to evaluate whether subjects used the same or different encoding mechanisms in each of the two environments. To answer this question, we tested which model, a generalized form of divisive normalization (DN) or power utility, better captures subjects’ choices. We picked this DN model because it is regarded as a canonical encoding mechanism in the brain (15–17, 19), including in the choice domain (7, 24, 25, 33). Crucially, the DN model has been considered an efficient encoder (1, 9, 13, 35), as at least one variant of the DN model has been proven to efficiently encode Pareto distributed environments (26). We thus expected some form of DN encoding in this environment. In the DN model, subject *i*’s STAGE II subjective value function of a lottery *k* ∈ {1,2} with payoffs *x*_1,*k*_ or *x*_2,*k*_ is given by:

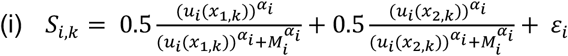

where *α*_*i*_ is the function’s curvature, *u*_*i*_(*·*) is subject *i*’s STAGE I valuation (*ρ*_*i*_), *M* is the reward expectation, and *ε*_*i*_ is an additive decision noise drawn in each trial from a zero-mean normal distribution, such that *ε*_*i*_∼*N*(0, *θ*_*DN*_).

The second model we examined was the commonly used power utility model (36):

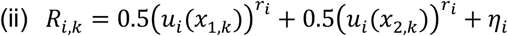

The model has one free parameter (*r*), which captures the function’s curvature. When *r* = 1, the function is linear. Similarly to our DN model, here, too, we included an additive decision noise *η*_*i*_∼*N*(0, *θ*_*p*_).

For every subject, we estimated both models using maximum likelihood estimation (see Materials and Methods). The subject-specific recovered parameters are reported in Table S4, and the sample medians are in Table 2. To determine, at the population level, which model better captured subjects’ choice patterns in each environment, we compared each subject’s Bayesian Information Criterion (BIC) scores across the two models in each environment^3^.

**Table 2.** Median estimates.

|  | Power Utility model |  |  | DN model |  |  |  |
| --- | --- | --- | --- | --- | --- | --- | --- |
| Parameter | r | $\theta_p$ | BIC | $\alpha$ | M | $\theta_{DN}$ | BIC |
| Uniform | 0.379 | 0.054 | 121.120 | 1.299 | 23.216 | 0.0233 | 104.670 |
| Pareto | 0.528 | 0.103 | 184.761 | 1.358 | 18.875 | 0.026 | 154.482 |

In line with our hypothesis, we found that in the Pareto environment, subjects’ BIC scores were on average significantly lower, indicating a better model fit, for the DN model than for the power utility model (Fig. 3C, one-sided Wilcoxon sign-rank test, Z=4.4603, p<0.0001). This was true for 48 subjects (out of 76). In the uniform environment, we expected that a model that does not belong to the family of DN models would better fit the data. Instead, we found that again the BIC scores were on average significantly lower for the DN model (Fig. 3D, one-sided Wilcoxon sign-rank test, Z=2.9692, p=0.0015) and this held for 42 (out of 76) subjects. Moreover, for only four subjects the curvature parameter in the power utility model was estimated as linear or as almost linear (*r* = 1 ± 0.05). The mean and median *r* estimates were 0.608 and 0.366, respectively. Importantly, the asymmetrical distributions of the differences in BIC scores (see insets in Fig. 3C-D) indicate that while for most subjects both models do (almost) equally well, there is a group of subjects for whom the DN model predicts their choices much better (ΔBIC>20 for 29 subjects in Pareto and 18 subjects in uniform).

To further compare the two models, we estimated them on the sample level. Table S5 presents the recovered pooled estimates from this analysis. Note that this analysis could only be done in dollar-space to allow comparability of lotteries across subjects, and to recover meaningful estimates of the M parameter in the DN model. Here, too, we find that the DN model captured subjects’ choices better, evident by the lower BIC scores when aggregating choices from both treatments (leftmost column), as well as within each environment (second and third columns). These results should be interpreted cautiously since the reward distributions were not fully controlled in the dollar space (Fig. S1).

Another way to examine the effect of the distributional environment on subjects’ value-encoding – and validate our task design – is to examine the relationship between the estimates of subjects’ subjective valuations of lotteries in STAGE I (*ρ*_*i*_, see Fig. 2E) and STAGE II α parameter in DN, and *r* parameter in power utility. Due to the nature of our design, a hyperbolic relationship (i.e.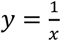) would suggest that STAGE I curvature was undone in STAGE II in both models. In the power utility model, this would also imply linear encoding of monetary payoffs^4^. In DN, it would mean that all curvature in STAGE II is associated with the DN encoding. Fig. 3E-G plots STAGE II parameters against STAGE I *ρ*.

In both Pareto and uniform environments, we saw a much closer hyperbolic relationship between *ρ* and the DN *α* parameter (Fig. 3 E and G) than between *ρ* and power utility *r* parameter (Fig. 3 F and H). We compared the root of mean squared errors (RMSE) between the hyperbolic function and the parameters in both models and confirmed that across the two environments, the α parameter of the DN model was more likely to maintain this hyperbolic relationship (α: Pareto: RMSE=0.6491, uniform: RMSE=0.6585; r: Pareto: RMSE=0.8109, uniform: RMSE=0.8588). This suggests that the curvature of the STAGE II subjective value functions can be attributed to DN value coding.

Taken together, all these results strengthen the notion that subjects used DN encoding of value in both environments.

### Context-Dependency: Adaptation of the Encoding Function to the Choice Environment

Our next aim was to examine whether subjects adapted their encoding according to the properties of the different environments. Another key difference between the uniform and Pareto environments was that, for all subjects in our sample, the medians of the subjective valuations in the uniform environments were higher than in Pareto (sample medians: 18.721 vs. 14.723 util units, respectively, Δ = 3.998, one-sided Wilcoxon sign-rank test between subject-specific medians, Z=7.572, p<0.0001). The reward expectation *M* in the DN model tracks the median of the reward distribution, and hence we hypothesized it would be higher in the uniform environment. Consistent with this hypothesis, the sample median of the recovered *M* parameters in the uniform environment was higher by 4.99 (in util units) than in the Pareto environment (one-sided Wilcoxon sign-rank test, Z=2.8907, p=0.0019, Table 1). This difference between the recovered *M* parameters was very close to the actual difference between the distributions’ medians, indicating that subjects – at least at the sample-level – quite precisely calibrated their encoding to the difference in reward expectation. On the subject level, we found that for 44 out of 76 subjects estimated *M* was higher in the uniform environment (Fig. 4A).

**Figure 4.**
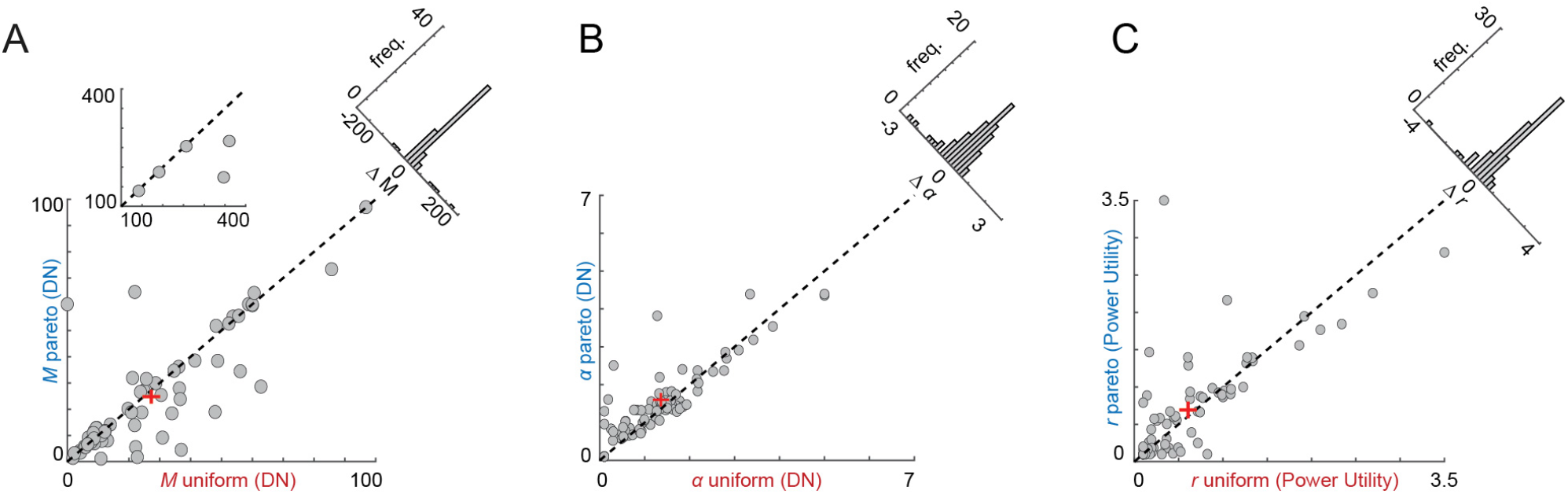
Cross-environment adaptation. (A-B) Adaptation of the encoding function in the DN model. (A) Best-fitting *M* parameter in the uniform (x-axis) vs. the Pareto (y-axis) environments. Estimates of *M*’s are in utility space. Left inset: outliers. Right (diagonal) inset: Difference in the estimates of *M* across choice environments (*M*(*uniform*) − *M*(*pareto*)). Insets do not show three additional (risk-seeking) subjects whose *M*’s are >400 (in util units). Dots indicate individual subjects, + indicate sample average without the inset outliers, N=76. (B) same as (A) for the DN’s *α* parameter. (C) Adaptation of the encoding function in the power utility model. Same as (B), but for the *r* parameter from the power utility model. (B-C) Dots indicate individual subjects, + indicate sample average, N=76.

Our pooled estimation further supports this conclusion with *M*(*uniform*)=66.6531 and *M*(*pareto*)=55.5609, second and third columns in Table S4, p<0.001. As a final robustness check, we estimated the DN model using the full dataset with the data from both environments, and included an additive dummy variable for the Pareto environment in the estimation of the *M* parameter (*M* = *constant* + *M*_pareto_*xpareto*). The output of this model split *M* into a constant, corresponding to the estimate of *M* for the uniform environment, and *M*_*pareto*_, which captured the difference in *M* in the Pareto relative to the uniform environment. We found *M*_*pareto*_ to be negative and significant (p<0.001), indicating *M* was lower in the Pareto environment.

In contrast to the *M* parameter, we had no prior hypotheses regarding the model’s curvature parameter α. Nevertheless, comparing subject-specific estimates, we found that on average, the α parameter was higher by 0.1593 in the Pareto environment (one-sided Wilcoxon sign-rank test, Z=1.9987, p=0.0228, Fig. 4B). This result may indicate that higher α values in the Pareto environment allowed better discriminability between the more frequently encountered lottery options, also indicated by our model-free analysis (Table 1), which was crucial given the correlational structure between the two valuations. However, this result was not fully replicated in the pooled estimates: when estimating each environment separately, we found that recovered parameters were almost identical (α_uniform_=0.93, α_Pareto_=0.92, Table S4, second and third columns), but a full model with random effect for the Pareto environment (similarly to the one run on *M*), revealed that there was a tuning of the function curvature when switching between environments (Table S4, rightmost column, p<0.001).

The power utility model is not designed to capture the dependence of the subjective value function on the distribution of valuations and hence we did not anticipate an adaptation of the function’s curvature. Indeed, when comparing estimates of *r* across the two environments, we obtain inconclusive results: while the pooled estimates indicated higher *r* values in the Pareto environment (Table S4), the subject-level estimates point in the opposite direction (Fig. 4C, one-sided Wilcoxon sign-rank test between subject-level estimates of *r*, Z=0.0511, p=0.4796).

To conclude this section, we found that subjects adapted the DN encoding function’s parameters to the two environments in line with our hypothesis, showing context-dependency in choice.

## Discussion

In this study, we tested how the distributional properties of choice environments affect value encoding. In particular, we were interested in whether the subjective value of rewards is encoded via a mechanism like divisive normalization (DN) exclusively in the Pareto environments for which it is efficient, or whether a DN representation is also employed in environments characterized by different reward distributions. To this end, we designed an experiment in which subjects were asked to make choices in two distinct statistical environments. In one environment, rewards were drawn from a Pareto distribution of valuations, for which DN is considered an efficient encoder. In the other environment, valuations were uniformly distributed, and DN would not represent the environment most efficiently (26).

Our results indicate that subjects in our study were better described as using a DN mechanism than a power utility mechanism to encode the subjective value of rewards, no matter from which of our two distributions the rewards were drawn. As expected, the key parameter of the model tracked the median of the distribution. A model-free analysis indicated that, compared to the uniform environment, when in the Pareto environment, subjects made fewer mistakes in choice sets drawn from the center of the distribution at the expense of the margins, a principal property of the DN function. We then fit our subjects’ choices with two classic stochastic choice models – one was a standard RUM with a power utility function, and the other was a RUM with a utility function belonging to the family of DN models. Our subject-level and pooled model-fitting results suggested that the DN model better captured subjects’ choice patterns in both the Pareto and the uniform environments (Table 2, Fig. 4C-D and Table S5). In line with the actual statistical properties of the two environments, subjects had higher reward expectations in the uniform environment. Taken together, these findings suggest that although subjects’ choices were affected by the context of the choice environment, their choice mechanisms were constrained to a DN encoding of value (Fig. 5).

**Figure 5.**
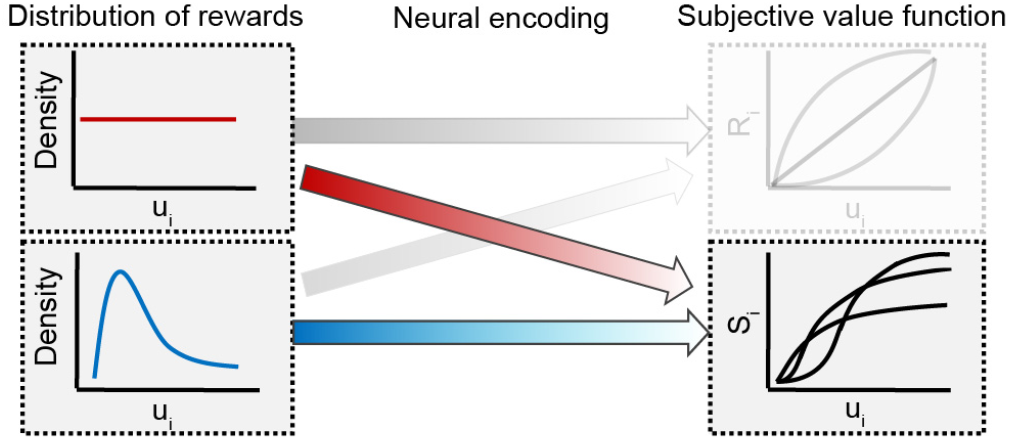
Summary of main findings.

Our findings indicate that in both environments the encoding of value was better described by a DN rather than a power utility encoder. Why would this be so? One possibility is that Pareto distributions are very common in the real world, and hence, the brain has evolved a constraint that accords well with natural environments. Indeed, numerous sensory stimuli are characterized by Pareto-like statistical properties (1, 14, 18, 27). On a larger scale, Pareto distributions also capture various ecological quantities, such as temporal and spatial measures of biodiversity (37–40). This is true also for environments that are related to value-based decisions because many economic and financial quantities in modern societies (41, 42), including consumption of several categories of consumer goods (43), have Pareto-like properties. An alternative hypothesis is that even the uniform distributions we examined are more efficiently dealt with by a DN encoder than the non-divisive power utility encoder. Indeed, recent theoretical advances (13) have shown that all encoders with limited precision or accuracy must incorporate an implicit divisive cost, a class of encoders of which DN is a member.

Another important finding is that, compared with the standard utility functions used in economics, DN provides the brain with an immensely flexible tool for the representation of choice options (32, 33). Given the specific parameterization we employed for DN, our model embeds the standard concave utility function, but is also suitable for capturing preferences that follow *S-shaped* functions, similar to the one suggested by Prospect Theory (44). DN further tracks the median of rewards (*expectations*), which allows for scale-invariant adjustments to different environments, while ensuring a fine discrimination between stimuli that are in the center of the distribution (2, 9, 13, 45). These adjustments – also evident in our data – give rise to context effects in choice processes (25, 33, 46, 47), and are also the core reason for some notable perceptual illusions (48, 49).

Our findings also imply that some choice patterns should not be regarded as built-in decision biases, errors, or mistakes. Rather, they reflect adjustments of the brain, as a constrained system, to its environment, thus reflecting a rational value-encoding mechanism (2, 13). Such an observation can explain the under-sampling of rare events when subjects adjust to new choice environments (50, 51) since the main focus of the system is on the mass of occurrences. On a broader view, our results could explain heterogeneity in individual decisions, such as the effect of one’s position along the long-tail distributions of socioeconomic measures (and their shapes) on the quality of healthcare (52), savings (53), and consumption (54) choices.

Finally, an interesting question that stems directly from our research is to what extent our results generalize beyond decision making processes to other cognitive functions, such as sensory processing. Even though various natural sensory stimuli are described by Pareto-like properties (14, 18, 45), we also frequently encounter, and are required to process, non-natural non-Pareto stimuli (55, 56). Our findings therefore invite further investigation into the effects of an obligate DN encoding on the sensory processing of non-Pareto stimuli.

## Materials and Methods

Some of the data in this manuscript have been used in the conference paper in reference (57).

### Experimental Design

*Valuation task (STAGE I)*. Our goal was to establish whether the brain employs different value encoding models in environments with different reward distributions. To eliminate any additional prior heterogeneity in subjects’ subjective valuations of money, we generated distributions of rewards in the subjective value (SV) space instead in dollar amounts (or expected values). To map the subject-specific SV space, we first recovered individual-specific subjective value functions over dollar amounts. To do this, in STAGE I, we used a valuation task, in which subjects reported their willingness to pay to participate in a lottery. See Table S1 for the list of 33 lotteries used in this task. On each trial, subjects were presented with a visualization of a 50-50 lottery on the computer screen and had to type in their willingness to pay to participate in it as a dollar amount (Figure 2A). For each lottery, the valuation could range between the current lottery’s minimal and maximal payoff, in $0.10 increments. All subjects completed the same 33 trials in an order randomized at the subject level. At the end of the session, the realization of one randomly selected trial was implemented for payment, using a Becker–DeGroot–Marschak (BDM) (58) procedure which was designed to elicit truthful valuations.

*Choice task (STAGE II)*. STAGE II was designed to test whether the distribution of rewards (lotteries with different subjective valuations) in a choice environment affects what value encoding model subjects use. Subjects were asked to choose the 50-50 lottery they preferred from two available options that varied from trial to trial. Lottery payoffs ranged between $0 and $60 in $0.10 increments. Overall, subjects made 640 binary choices that were divided into two blocks of 320 trials each and presented on subsequent days. Our experimental manipulation was that in each block, the valuations were drawn either from a Pareto Type III distribution for which DN is an efficient code (26) or from a uniform distribution (Fig. 2A-B). The order in which subjects experienced these environments was counter-balanced across subjects. One trial was randomly selected for payment at the end of each experimental session.^5^

*Subjective Value of Money*. We used each subject’s STAGE I single lottery valuations to estimate their subjective value function over money. We expressed each subject *i*’s subjective value of a 50-50 lottery that paid *y*_1_ or *y*_2_, each equally likely, using a power utility function as:

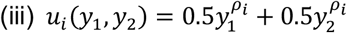

If the curvature parameter *ρ*_*i*_ < 1, then subject *i* is risk-averse. When *ρ*_*i*_ = 1, the subject is risk-neutral. If *ρ*_*i*_ > 1, the subject is risk-seeking. Therefore, the certainty equivalents (*c*) that participants stated were converted to subjective values using the same power utility function such that *c* = *u*^1/ ρ^. We ran an NLS regression to estimate the *ρ* parameter separately for each subject.

We used the subject’s estimated *ρ*_*i*_, to pick different combinations of lottery dollar payoffs to create lotteries that had a specific SV to that individual. This enabled us to generate sets of lotteries whose implied SV distributions matched our target distributions (see below), regardless of individual differences in the curvature of the subjective value function.

### Distributions of Valuations

*Uniform Distributions of SVs*. For each subject *i*, we computed the upper bound of the distribution as the SV of the maximal possible monetary payoff in the study, which was $60 (i.e.,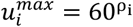). We then divided the range [0, 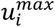] into 40 equally-spaced SV increments. For each of the increments, we created eight different lotteries, which would give the subject the subjective value in exactly this bracket (for a total of 320 lotteries). Since the joint distribution of a two-dimensional uniform distribution is independent, and hence determined by its marginals, we then picked pairs of lotteries from this set for generating binary choice sets.

*Pareto Type III Distributions of SVs*. The DN encoding function is information-maximizing for a bivariate Pareto distribution with a joint pdf (see Eq. 7 with *μ*_1_ = 0 in reference (26))^6^ *f*_*ui*_ (*u*_*i*,1_, *u*_*i*,2_) for every subject *i* and *k* ∈ {1,2} is an index indicating the choice option within the choice set.

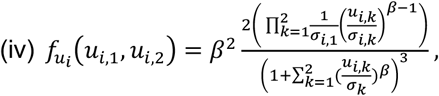

and the marginal pdf being a univariate Pareto Type III pdf:

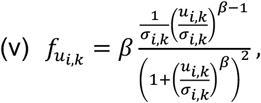

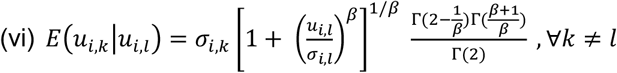

We matched, for each subject, *E*(*u*_*i*,*k* |_ *u*_*i*,*l*_) to the expectation of the uniform distribution, which was $30 (and ū = 30^ρ^ in SV-space), where Γ indicates the gamma function.

Following Proposition 4 in (26) and using the subject-specific parameterization, we generated the Pareto Type III distributions as a scale mixture of transformed exponential (or Weibull) random variables, so that:

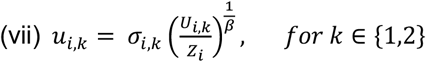

where *U*_*k*_∼*Exp*(λ = 1) and *Z*∼*Exp*(λ = 1) independently of all *U*_*k*_. Fig. 2C presents three examples for such distributions with different *ρ*_*i*_ values.

Note that using only 320 draws may lead to under-sampling of the distributions. Therefore, to fully capture the shape of the distribution, for each subject, we first generated joint Pareto distributions with 100K draws. We then created small 600-draw experimental distributions that matched the large 100k-draw distributions, allowing a deviation of up to 0.2 utils from the actual first and second moments (mean and standard deviation) of the large 100k-draws sets. Fig. 2D compares matched and unmatched small sets, corresponding to the large 100k-draws set presented in Fig. 2C (middle panel). Finally, we truncated the long tail of the Pareto Type III distributions at 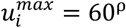 (eliminating 6.5 to 23.83 percent of the distribution, depending on the *ρ* parameter, the curvature of the subjective value function), to match the upper bound of the uniform distribution and to avoid extreme reward amounts. We then casted 320 SVs at random from the remaining valuations, which constituted the experimental subject-specific Pareto distributions.

*Generating Binary Choice Sets from the Distributions of Valuations*. The final step was to generate lottery dollar amounts from the SV distributions. For each lottery *k* with a valuation (*u*_*k*_), we first randomly drew the first monetary payoff *x*_1,*k*_ from a range of possible payoffs $0 − *x*^*max*^ in $0.10 increments. We had to restrict the maximum value of *x*_1,*k*_ to make sure that including it in the lottery, does not exceed the lottery valuation (*u*_*k*_), and thus to avoid negative values for the second lottery payoff. We determined the maximal value of the first payoff *x*_1,*k*_ using the minimum function:

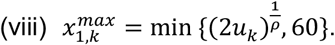

We then solved for *x*_*k*,2_, giving rise to the desired *u*_*k*_, rounded to one decimal place, using the following equation:

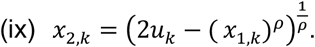

Figure S1 shows how the heterogeneity in ρ values affected the distributions of *x*_*k*,1_ and *x*_*k*,2_.

We restricted the share of trials with first-order stochastic dominance (FOSD) (trials on which both lottery payoffs of one lottery were higher or equal to the other lottery’s payoffs) to 45 percent. For subjects with ρ_*i*_ → 0, we could not generate experimental sets with only 45 percent of the trials. Thus, we fixed ρ_*i*_=1, for all subjects with ρ_*i*_ < 0.1 (a total of 4 subjects, see Table S2), limiting the interoperability of data from this small number of subjects. In contrast, for two subjects with very high ρ’s (ρ_i_ > 4), we also had to fix ρ_i_ = 1 in STAGE II of the study, since a very large tail from their Pareto distribution of SVs exceeded $60. Respectively, the interoperability of data from this subject is also limited. Nonetheless, we wanted to avoid any unjustified elimination of data, and therefore analyzed data from these six subjects. Importantly, our main qualitative findings do not change once we remove these subjects from our sample.

### Procedures

#### Sessions

Experimental sessions were carried out online via Zoom while subjects completed the task on a website. We ran eight sessions of the experiment between May 2022 and August 2022. After instruction, subjects had to successfully answer a set of comprehension questions about the procedure before starting STAGE I. They could participate in STAGE II of the study only if they completed all trials in STAGE I. Subjects received all payments after completing both STAGE I and STAGE II. Subjects received a $10 participation fee and on average $24.5 in STAGE I (range $0-60) and $76.02 in STAGE II (range $7.3-120) from the decision task. All amounts are in Australian dollars. All parts of the experiment were self-paced. Both the valuation and the choice tasks were programmed in the oTree software package (60).

#### Participants

We recruited participants from various departments at the University of Sydney. Subjects gave informed written consent before participating in the study, which was approved by the local ethics committee at the University of Sydney. Seventy-six subjects (44 females, mean age=21.8, std: 3.34, range: 18-30) passed the comprehension questions and completed STAGE I and the two choice tasks of STAGE II.

### Model Fitting

#### Sample-level (pooled) estimates

We estimated subjects’ aggregated choice data via a probit choice function with maximum likelihood estimation (MLE). Standard errors were clustered at the subject level. Thus, in the pooled estimation subjects were treated as one representative decision-maker. In this analysis we used lotteries’ monetary rewards (as opposed to their subjective valuations) to allow meaningful estimates of DN’s *M* parameter, and to confine the range of lottery payoffs. For both DN and power utility, we report the results from models estimated on the full dataset and separately on each choice environment.

To test the possibility of adaptation of the encoding function to the choice environments, we further report the results from three additional models estimated on the full dataset, which also included a dummy variable indicating the Pareto environment for the reward expectation, *M* parameter (DN) as *M* = *constant M*_*pareto*_*xpareto* and similarly for the functions’ curvature parameters *α* (DN) and *r* (power utility), respectively.

#### Subject-level estimates

*DN*. In each choice environment, we recovered subject-specific estimates of the free parameters, restricting the search space as follows: 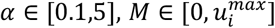 and *θ*>0 (see equation (i) in the text). We employed MLE using the Nelder-Mead algorithm with a max-iteration limit of 1,000 and a stopping criteria of 0.5 tolerance. We initialized *M* to the distributions’ medians. *θ* was initialized at 0.03, matching the sample-level pooled estimate (see Table S5). For the *α* parameter, we took ten random initializations in the range {0.1,5} with a precision of 5. For calculating the likelihoods, in each of the 320 trials, we generated 10,000 samples with randomly drawn Gaussian noise. The log-likelihood function was thus given by –

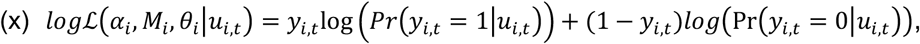

where *y*_*i*,*t*_ = {0,1} indicates the subject’s *i* choice in trial *t* = {1, ߪ, 320}.

*Power utility*. We fitted the power utility model to recover subject-specific estimates of the *r* and *θ* parameters using a similar procedure. We restricted the search space as follows: *r* ∈ {0.1,5}, and *θ*>0 (see equation (ii) in the text). *θ* was initialized at 0.03, matching the sample-level pooled estimate (see Table S5). For the *r* parameter, we took ten random initializations in the range {0.1,5} with a precision of 5. All other procedures were identical to the DN model.

## Supporting Information

**Figure S1.**
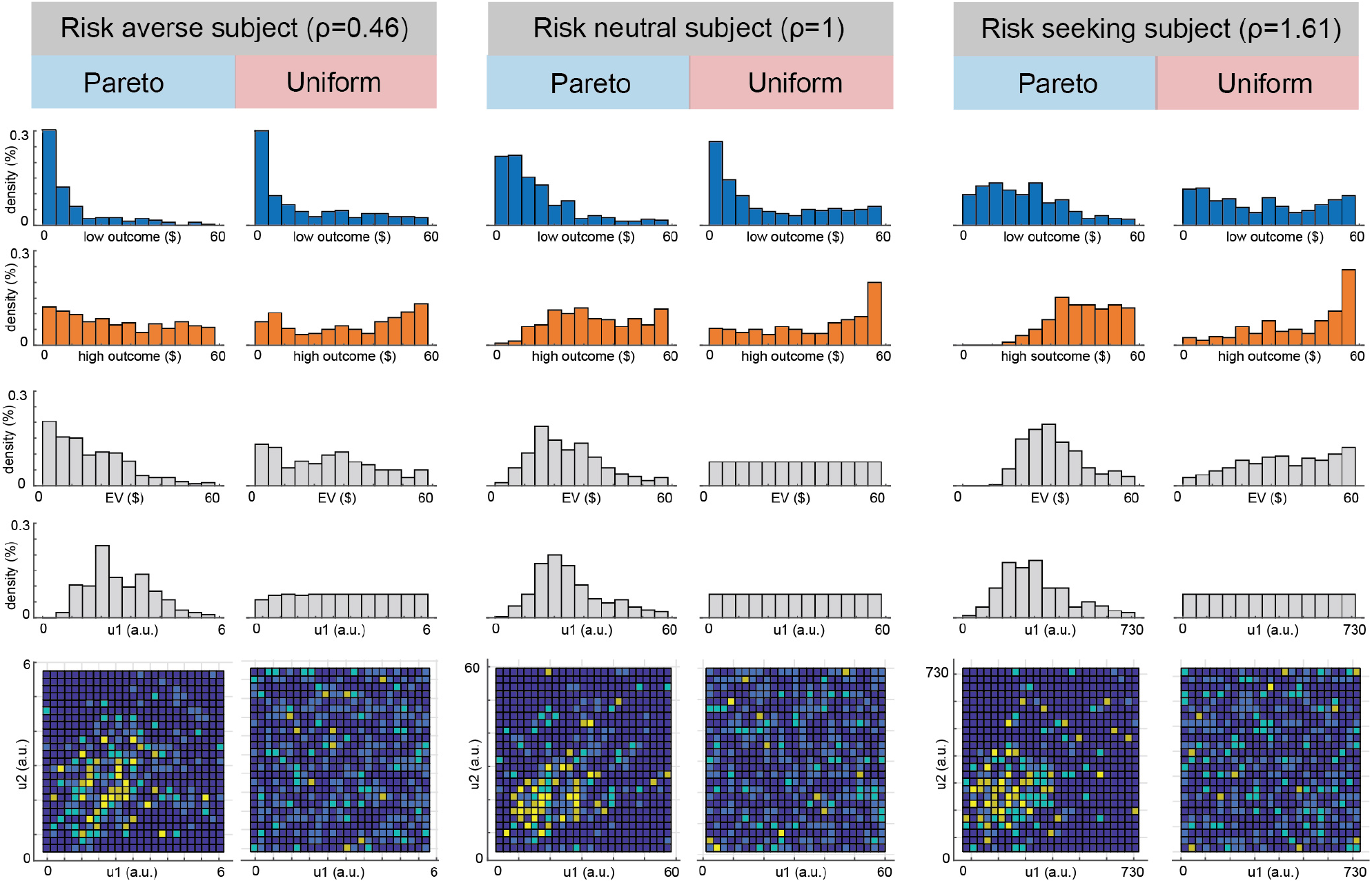
Representative Sets in STAGE II. Left – a risk averse subject, middle – a risk neutral subject, right – a risk seeking subject. Top to bottom: (1) Distributions of the high winning amount in Lottery 1 (in dollars); (2) Distributions of the low winning amount in lottery 1 (in dollars); (3) Distribution of the expected earnings (EV) of Lottery 1 (in dollars); (4) Distributions of the valuations (u1) of Lottery 1 (in util units); (5) 2-dimensional histogram of the valuations of Lottery 1 and Lottery 2 (u1 and u2, in util units).

**Figure S2.**
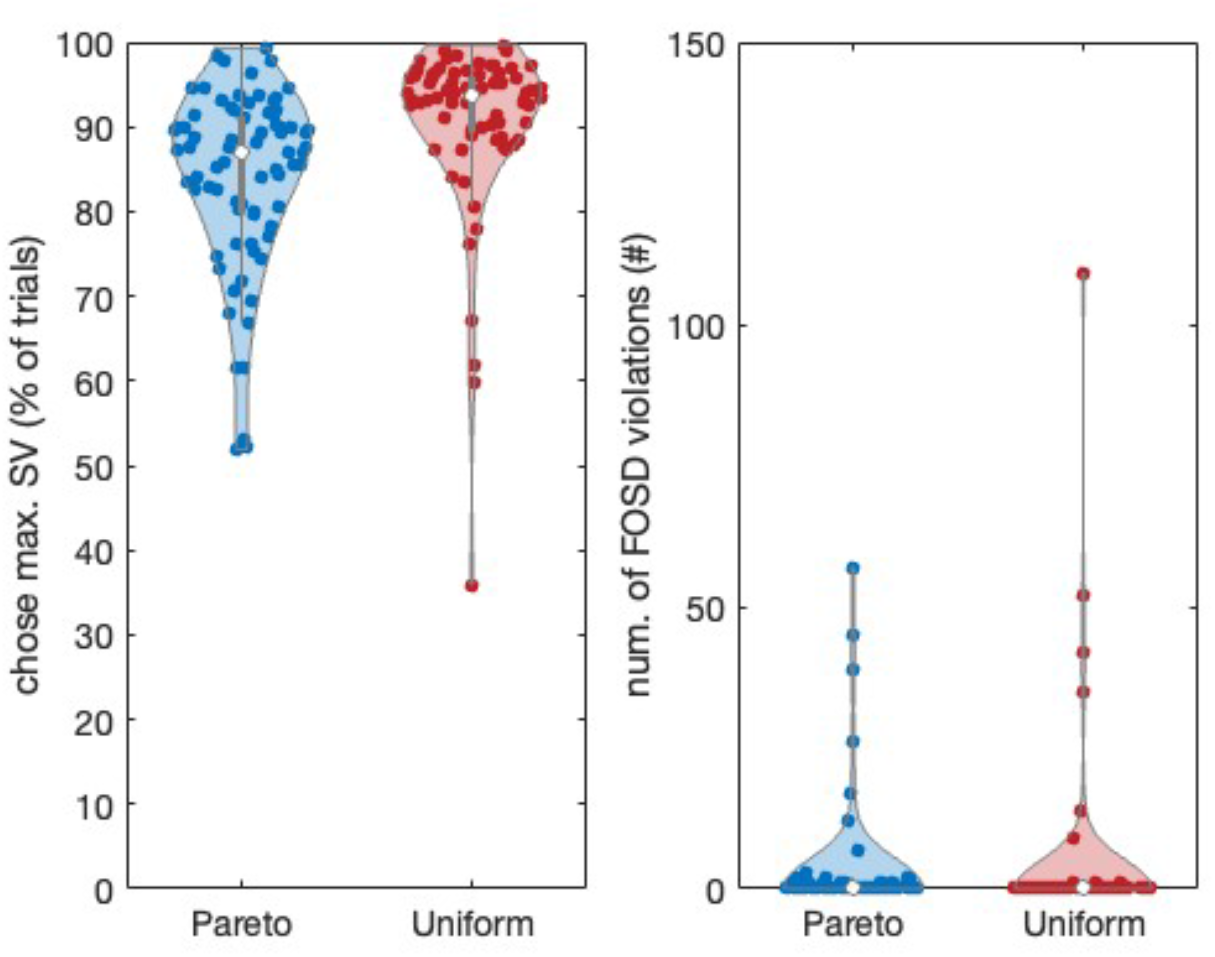
Descriptive statistics. Left – violins show the share of trials in which subjects chose the lottery with the higher subjective value. Right – violins show the number of FOSD violations per subject. Dots indicate individual subjects. N=76.

**Figure S3.**
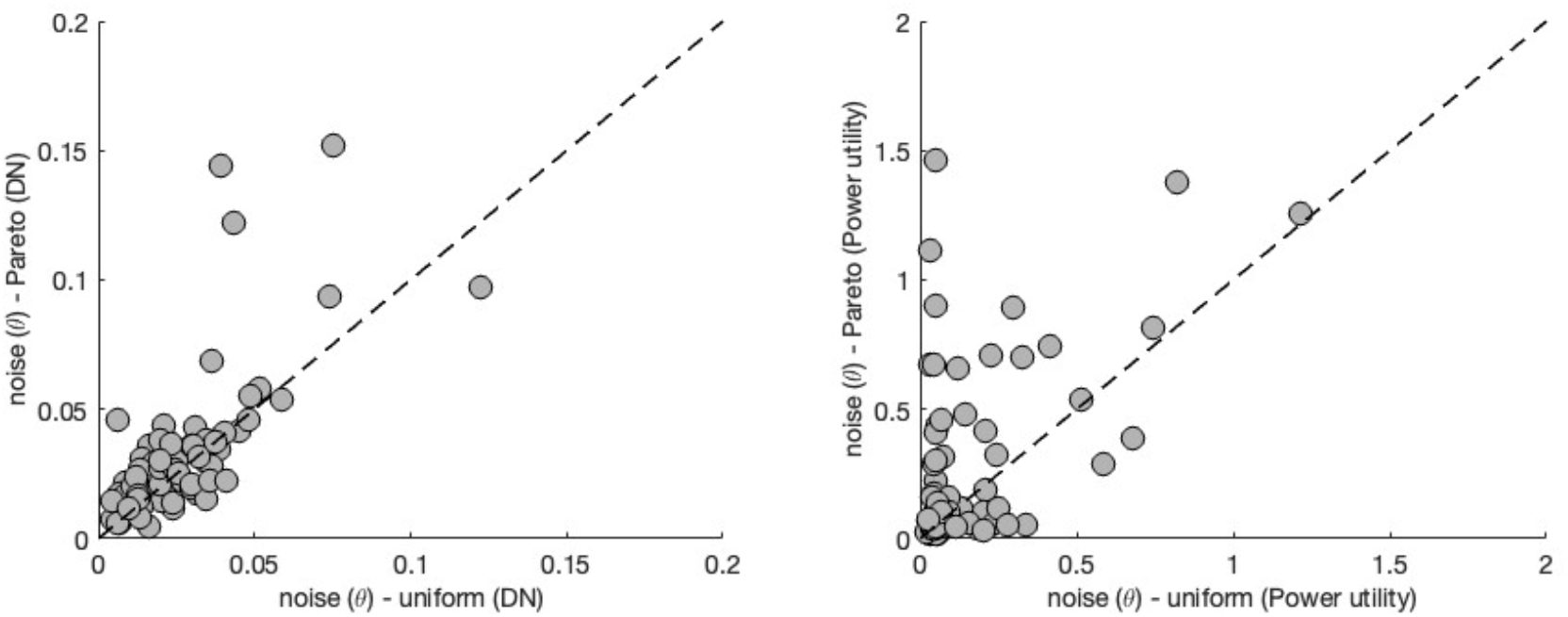
Noise estimates. Comparing the best-fitting *σ* parameter (decision noise) across the distributional environments reveals noise levels were higher in the Pareto environment. *Left* - DN model (one-sided Wilcoxon sign-rank test, Z=2.2314, p=0.0257). *Right* - Powe Utility model (one-sided Wilcoxon sign-rank test, Z=2.9172, p=0.0035). Scatters indicate individual subjects. N=76.

**Table S1.** Lotteries used in STAGE I

| lottery | x1 | x2 |
| --- | --- | --- |
| 1 | 60 | 0 |
| 2 | 55 | 0 |
| 3 | 50 | 0 |
| 4 | 45 | 0 |
| 5 | 40 | 0 |
| 6 | 35 | 0 |
| 7 | 30 | 0 |
| 8 | 25 | 0 |
| 9 | 20 | 0 |
| 10 | 15 | 0 |
| 11 | 10 | 0 |
| 12 | 5 | 0 |
| 13 | 60 | 5 |
| 14 | 55 | 5 |
| 15 | 50 | 5 |
| 16 | 45 | 5 |
| 17 | 40 | 5 |
| 18 | 35 | 5 |
| 19 | 30 | 5 |
| 20 | 25 | 5 |
| 21 | 20 | 5 |
| 22 | 15 | 5 |
| 23 | 10 | 5 |
| 24 | 60 | 10 |
| 25 | 55 | 10 |
| 26 | 50 | 10 |
| 27 | 45 | 10 |
| 28 | 40 | 10 |
| 29 | 35 | 10 |
| 30 | 30 | 10 |
| 31 | 25 | 10 |
| 32 | 20 | 10 |
| 33 | 15 | 10 |

**Table S2.** Individual-level estimates of risk preferences from subjects’ bids in STAGE I.

| SID | $\rho$ | SE | SID | $\rho$ | SE |
| --- | --- | --- | --- | --- | --- |
| 1005 | 0.000 (*) | 0.009 | 1501 | 0.866 | 0.051 |
| 1104 | 0.000 (*) | 0 | 1406 | 0.875 | 0.018 |
| 1205 | 0.000 (*) | 0 | 1308 | 0.876 | 0.093 |
| 1401 | 0.000(*) | 0 | 1614 | 0.876 | 0.068 |
| 1001 | 0.347 | 0.055 | 908 | 0.88 | 0.045 |
| 1006 | 0.374 | 0.067 | 1508 | 0.891 | 0.066 |
| 1301 | 0.377 | 0.056 | 1408 | 0.909 | 0.033 |
| 1201 | 0.429 | 0.106 | 909 | 0.927 | 0.176 |
| 1617 | 0.437 | 0.04 | 1008 | 0.946 | 0.055 |
| 903 | 0.443 | 0.045 | 1502 | 0.954 | 0.065 |
| 1619 | 0.461 | 0.036 | 1607 | 0.954 | 0.042 |
| 1605 | 0.463 | 0.094 | 1402 | 0.961 | 0.067 |
| 1601 | 0.49 | 0.057 | 1604 | 0.968 | 0.018 |
| 1101 | 0.492 | 0.069 | 1105 | 0.981 | 0.018 |
| 910 | 0.499 | 0.053 | 1204 | 1 | 0 |
| 1610 | 0.5 | 0.043 | 1603 | 1 | 0 |
| 1405 | 0.518 | 0.048 | 1409 | 1.008 | 0.132 |
| 906 | 0.586 | 0.039 | 1513 | 1.012 | 0.058 |
| 1407 | 0.593 | 0.045 | 1012 | 1.018 | 0.028 |
| 1611 | 0.601 | 0.054 | 1613 | 1.018 | 0.016 |
| 1106 | 0.612 | 0.058 | 1505 | 1.047 | 0.042 |
| 1109 | 0.627 | 0.044 | 1608 | 1.09 | 0.157 |
| 1307 | 0.628 | 0.039 | 1305 | 1.185 | 0.126 |
| 1403 | 0.639 | 0.06 | 1004 | 1.192 | 0.145 |
| 1615 | 0.64 | 0.084 | 1002 | 1.201 | 0.068 |
| 904 | 0.647 | 0.048 | 1504 | 1.203 | 0.142 |
| 1510 | 0.655 | 0.074 | 1102 | 1.244 | 0.086 |
| 902 | 0.686 | 0.047 | 1003 | 1.253 | 0.062 |
| 1107 | 0.711 | 0.227 | 1609 | 1.278 | 0.095 |
| 1303 | 0.718 | 0.038 | 1304 | 1.358 | 0.197 |
| 1103 | 0.727 | 0.038 | 1503 | 1.407 | 0.109 |
| 1010 | 0.733 | 0.041 | 1302 | 1.577 | 0.143 |
| 1512 | 0.752 | 0.025 | 1011 | 1.61 | 0.157 |
| 1506 | 0.769 | 0.036 | 1616 | 1.808 | 0.24 |
| 1507 | 0.775 | 0.074 | 912 | 2.276 | 0.185 |
| 1306 | 0.806 | 0.055 | 1202 | 124.189 (*) | 28.955 |
| 907 | 0.819 | 0.045 | 1203 | 4.432 (*) | 0.261 |
| 1014 | 0.833 | 0.026 |  |  |  |

(\*) For these subjects we could not generate distributions of valuations for STAGE II that would adhere to our requirement to limit the number of trials with FOSD violations (when $\rho_i \rightarrow 0$ ), or without having to censor a very large tail of the Pareto distribution (when $\rho_i > 4$ ). Instead, for these subjects we plugged-in $\rho_i = 1$ to generate the distributions for STAGE II.

**Table S3.** Robustness checks for the findings presented in Column (2) in Table 1. We vary the definitions for *center of the distributions* (center) and *around the diagonal* (diagonal). Column (1) corresponds to the regression presented in the Main Text.

| | (1)<br>Center: \$9-42<br>Diagonal: $0.9 < \frac{u_2}{u_1} < 1.1$ | (2)<br>Center: \$9-42<br>Diagonal: $0.95 < \frac{u_2}{u_1} < 1.05$ | (3)<br>Center: \$12-39<br>Diagonal: $0.9 < \frac{u_2}{u_1} < 1.1$ | (4)<br>Center: \$12-39<br>Diagonal: $0.95 < \frac{u_2}{u_1} < 1.05$ |
| --- | --- | --- | --- | --- |
| SV difference | -0.0001<br>(0.0000) | -0.0000<br>(0.0000) | -0.0002**<br>(0.0001) | -0.0001**<br>(0.0000) |
| Pareto | -0.0003***<br>(0.0001) | -0.0003***<br>(0.0001) | -0.0002***<br>(0.0001) | -0.0002**<br>(0.0001) |
| Pareto*SV difference | -0.1479***<br>(0.0374) | -0.1547***<br>(0.0334) | -0.1138**<br>(0.0394) | -0.1228***<br>(0.0354) |
| Near diagonal | -0.7400***<br>(0.0692) | -0.8916***<br>(0.0730) | -0.6463***<br>(0.0708) | -0.8166***<br>(0.0740) |
| Pareto*Near diagonal | 0.1742**<br>(0.0597) | 0.3213***<br>(0.0660) | 0.1427*<br>(0.0656) | 0.3178***<br>(0.0734) |
| Constant | 1.1201***<br>(0.0713)<br>22442 | 1.0618***<br>(0.0635)<br>22442 | 1.0146***<br>(0.0726)<br>16576 | 0.9580***<br>(0.0642)<br>16576 |
| N | 0.036 | 0.027 | 0.029 | 0.022 |
| pseudo R-sq | -0.0001 | -0.0000 | -0.0002** | -0.0001** |

**Table S4.** Individual-level best-fitting model parameters across environments (STAGE II).

| SID | $\rho$ | DN | | | | | | Power Utility | | | |
| --- | --- | --- | --- | --- | --- | --- | --- | --- | --- | --- | --- |
|  |  | Pareto |  |  | Uniform |  |  | Pareto |  | Uniform |  |
| | | M | $\alpha$ | $\theta$ | M | $\alpha$ | $\theta$ | r | $\theta$ | r | $\theta$ |
| 1001 | 0.35 | 4.063 | 3.54 | 0.031 | 4.142 | 3.851 | 0.014 | 2.258 | 1.257 | 2.692 | 1.215 |
| 1006 | 0.37 | 1.442 | 2.4 | 0.058 | 1.808 | 1.838 | 0.051 | 0.674 | 0.192 | 0.714 | 0.207 |
| 1301 | 0.38 | 2.713 | 4.39 | 0.122 | 2.52 | 3.342 | 0.043 | 1.319 | 0.747 | 1.265 | 0.413 |
| 1201 | 0.43 | 5.709 | 2.7 | 0.025 | 5.517 | 2.82 | 0.02 | 1.947 | 1.376 | 1.918 | 0.819 |
| 903 | 0.44 | 3.642 | 3.18 | 0.043 | 3.622 | 3.417 | 0.031 | 1.347 | 0.539 | 1.336 | 0.517 |
| 1617 | 0.44 | 5.583 | 2.05 | 0.03 | 5.938 | 2.195 | 0.02 | 1.397 | 0.711 | 1.274 | 0.227 |
| 1619 | 0.46 | 6.615 | 2.86 | 0.012 | 6.615 | 2.795 | 0.01 | 1.767 | 0.456 | 2.1 | 0.067 |
| 1101 | 0.49 | 7.491 | 4.39 | 0.014 | 7.491 | 5 | 0.024 | 2.803 | 0.032 | 3.5 | 50.499 |
| 1601 | 0.49 | 3.221 | 1.29 | 0.026 | 2.299 | 1.713 | 0.024 | 0.608 | 0.142 | 0.463 | 0.054 |
| 910 | 0.5 | 7.726 | 2.91 | 0.012 | 7.726 | 3.091 | 0.024 | 1.843 | 0.067 | 2.341 | 0.028 |
| 1610 | 0.5 | 7.439 | 2.14 | 0.046 | 7.732 | 2.158 | 0.048 | 1.466 | 1.463 | 0.169 | 0.051 |
| 1405 | 0.52 | 8.329 | 2.35 | 0.007 | 8.337 | 2.52 | 0.005 | 1.558 | 0.046 | 1.862 | 0.035 |
| 906 | 0.59 | 9.181 | 1.48 | 0.038 | 6.865 | 1.268 | 0.035 | 0.84 | 0.418 | 0.628 | 0.21 |
| 1407 | 0.59 | 10.038 | 2.19 | 0.032 | 11.066 | 1.331 | 0.032 | 1.397 | 0.041 | 0.605 | 0.187 |
| 1611 | 0.6 | 10.916 | 1.47 | 0.021 | 8.483 | 1.932 | 0.03 | 0.971 | 0.391 | 1.092 | 0.678 |
| 1106 | 0.61 | 11.711 | 3.82 | 0.069 | 12.268 | 1.277 | 0.037 | 3.5 | 0.016 | 0.335 | 0.054 |
| 1109 | 0.63 | 7.008 | 1.82 | 0.025 | 8.8 | 1.596 | 0.02 | 0.856 | 0.324 | 0.774 | 0.242 |
| 1307 | 0.63 | 12.895 | 1.8 | 0.021 | 9.504 | 1.48 | 0.028 | 1.29 | 0.9 | 0.61 | 0.05 |
| 1403 | 0.64 | 10.037 | 2.37 | 0.008 | 9.728 | 2.764 | 0.013 | 1.392 | 0.818 | 1.331 | 0.744 |
| 1615 | 0.64 | 8.884 | 0.71 | 0.023 | 8.685 | 0.911 | 0.041 | 0.231 | 0.048 | 0.348 | 0.114 |
| 904(**) | 0.65 | 14.149 | 0.1 | 1 | 14.149 | 0.1 | 1 | 0.1 | 3.84 | 0.1 | 1.174 |
| 1510 | 0.66 | 12.807 | 1.06 | 0.02 | 9.392 | 1.328 | 0.03 | 0.599 | 0.162 | 0.444 | 0.041 |
| 902 | 0.69 | 8.798 | 1.57 | 0.033 | 8.454 | 1.776 | 0.018 | 0.249 | 0.051 | 0.717 | 0.224 |
| 1107 | 0.71 | 3.503 | 1.59 | 0.042 | 4.333 | 1.461 | 0.045 | 0.294 | 0.111 | 0.193 | 0.053 |
| 1303 | 0.72 | 8.09 | 1.55 | 0.027 | 13.044 | 1.146 | 0.018 | 0.556 | 0.158 | 0.479 | 0.093 |
| 1010 | 0.73 | 20.129 | 1.33 | 0.036 | 19.703 | 1.466 | 0.021 | 0.874 | 0.89 | 0.747 | 0.296 |
| 1103 | 0.73 | 7.997 | 0.79 | 0.03 | 11.232 | 0.562 | 0.025 | 0.255 | 0.073 | 0.162 | 0.044 |
| 1512 | 0.75 | 11.392 | 1.76 | 0.054 | 12.119 | 1.32 | 0.059 | 0.165 | 0.05 | 0.154 | 0.054 |
| 1506 | 0.77 | 18.808 | 1.55 | 0.024 | 20.815 | 1.373 | 0.012 | 0.935 | 0.676 | 0.63 | 0.045 |
| 1507(**) | 0.77 | 1.15 | 2.51 | 0.055 | 10.924 | 0.306 | 0.049 | 0.1 | 0.101 | 0.1 | 0.09 |
| 1306 | 0.81 | 27.078 | 1.83 | 0.021 | 24.896 | 2.176 | 0.02 | 0.886 | 0.437 | 0.143 | 0.054 |
| 907 | 0.82 | 11.66 | 1.15 | 0.036 | 8.719 | 1.231 | 0.016 | 0.181 | 0.035 | 0.391 | 0.07 |
| 1014 | 0.83 | 18.769 | 1.42 | 0.03 | 24.162 | 1.355 | 0.034 | 0.663 | 0.288 | 0.742 | 0.587 |
| 1501 | 0.87 | 34.674 | 0.82 | 0.023 | 34.617 | 0.883 | 0.027 | 0.183 | 0.049 | 0.196 | 0.049 |
| 908 | 0.88 | 27.922 | 0.99 | 0.021 | 36.342 | 0.95 | 0.018 | 0.55 | 0.227 | 0.255 | 0.047 |
| 1308 | 0.88 | 2.713 | 4.39 | 0.122 | 36.133 | 2.176 | 0.03 | 0.1 | 0.061 | 0.1 | 0.033 |
| 1406 | 0.88 | 29.879 | 1.57 | 0.016 | 28.679 | 1.578 | 0.008 | 0.923 | 0.674 | 0.952 | 0.03 |
| 1614 | 0.88 | 31.894 | 1.31 | 0.022 | 21.161 | 1.267 | 0.036 | 0.2 | 0.074 | 0.1 | 0.026 |
| 1508 | 0.89 | 1.747 | 0.87 | 0.041 | 22.703 | 0.518 | 0.041 | 0.1 | 0.04 | 0.1 | 0.04 |
| 1408 | 0.91 | 38.529 | 1.34 | 0.015 | 41.378 | 1.638 | 0.004 | 0.865 | 0.656 | 1.001 | 0.121 |
| 909 | 0.93 | 18.397 | 0.66 | 0.035 | 33.938 | 0.65 | 0.039 | 0.156 | 0.052 | 0.114 | 0.044 |
| 1008 | 0.95 | 13.914 | 1.33 | 0.044 | 21.842 | 0.822 | 0.021 | 0.183 | 0.054 | 0.219 | 0.05 |
| 1502 | 0.95 | 18.941 | 0.52 | 0.021 | 47.922 | 0.293 | 0.011 | 0.1 | 0.022 | 0.1 | 0.022 |
| 1607 | 0.95 | 31.53 | 1.38 | 0.038 | 25.643 | 1.5 | 0.038 | 0.133 | 0.055 | 0.523 | 0.278 |
| 1402 | 0.96 | 25.401 | 1.35 | 0.036 | 30.41 | 1.333 | 0.031 | 0.575 | 0.3 | 0.225 | 0.053 |
| 1604 | 0.97 | 52.639 | 1.37 | 0.006 | 52.397 | 1.779 | 0.006 | 0.906 | 0.042 | 1.087 | 0.043 |
| 1105 | 0.98 | 4.505 | 1.3 | 0.046 | 36.939 | 0.1 | 0.006 | 0.1 | 0.03 | 0.1 | 0.038 |
|  |  | Pareto |  |  | Uniform |  |  | Pareto |  | Uniform |  |
| | | M | $\alpha$ | $\theta$ | M | $\alpha$ | $\theta$ | r | $\theta$ | r | $\theta$ |
| 1005 | 1(*) | 55.271 | 1.04 | 0.017 | 53.736 | 0.573 | 0.007 | 0.582 | 0.286 | 0.29 | 0.045 |
| 1013 | 1(*) | 60 | 1.54 | 0.004 | 58.779 | 1.346 | 0.016 | 0.984 | 0.319 | 0.881 | 0.075 |
| 1104 | 1(*) | 5.542 | 0.83 | 0.015 | 22.165 | 0.247 | 0.009 | 0.1 | 0.014 | 0.134 | 0.033 |
| 1202 | 1(*) | 59.606 | 1.44 | 0.015 | 60 | 1.698 | 0.009 | 0.999 | 1.115 | 1.229 | 0.03 |
| 1203(**) | 1(*) | 60 | 0.1 | 1 | 60 | 0.1 | 1 | 0.1 | 5.389 | 0.1 | 42.832 |
| 1204 | 1 | 59.894 | 1.46 | 0.006 | 60 | 1.603 | 0.007 | 0.947 | 0.041 | 1.016 | 0.034 |
| 1205 | 1(*) | 9.181 | 1.02 | 0.028 | 30.88 | 0.569 | 0.036 | 0.1 | 0.021 | 0.127 | 0.05 |
| 1401 | 1(*) | 34.454 | 0.77 | 0.015 | 56.078 | 0.928 | 0.012 | 0.305 | 0.071 | 0.364 | 0.053 |
| 1603 | 1 | 55.57 | 1.42 | 0.015 | 55.573 | 1.85 | 0.035 | 0.896 | 1.05 | 0.991 | 2.048 |
| 1409 | 1.01 | 23.842 | 0.96 | 0.144 | 36.602 | 0.1 | 0.039 | 0.1 | 0.106 | 0.1 | 0.204 |
| 1513 | 1.01 | 28.602 | 0.94 | 0.017 | 62.8 | 1.125 | 0.013 | 0.391 | 0.117 | 0.645 | 0.247 |
| 1012 | 1.02 | 64.681 | 0.76 | 0.018 | 21.882 | 0.885 | 0.025 | 0.431 | 0.175 | 0.191 | 0.047 |
| 1613 | 1.02 | 64.294 | 1.08 | 0.015 | 60.575 | 0.993 | 0.013 | 0.672 | 0.412 | 0.404 | 0.048 |
| 1505 | 1.05 | 26.584 | 1.46 | 0.027 | 23.729 | 1.501 | 0.02 | 0.187 | 0.059 | 0.495 | 0.236 |
| 1608 | 1.09 | 73.339 | 1.74 | 0.093 | 85.689 | 1.691 | 0.074 | 0.1 | 0.101 | 0.1 | 0.066 |
| 1305(**) | 1.18 | 97.004 | 0.1 | 0.152 | 96.82 | 0.1 | 0.075 | 0.1 | 0.705 | 0.1 | 0.326 |
| 1004 | 1.19 | 103.965 | 0.693 | 0.013 | 59.74 | 0.796 | 0.026 | 0.352 | 0.135 | 0.278 | 0.116 |
| 1002 | 1.2 | 38.408 | 1.35 | 0.022 | 48.763 | 1.09 | 0.012 | 0.179 | 0.051 | 0.353 | 0.083 |
| 1504 | 1.2 | 137.745 | 0.76 | 0.027 | 137.959 | 0.299 | 0.013 | 0.1 | 0.041 | 0.177 | 0.048 |
| 1102 | 1.24 | 51.763 | 1.32 | 0.038 | 48.245 | 0.921 | 0.02 | 0.124 | 0.047 | 0.2 | 0.044 |
| 1003 | 1.25 | 40.119 | 0.62 | 0.015 | 143.863 | 0.527 | 0.02 | 0.179 | 0.04 | 0.159 | 0.049 |
| 1609 | 1.28 | 186.451 | 1.05 | 0.026 | 187.422 | 0.995 | 0.026 | 0.1 | 0.063 | 0.311 | 0.154 |
| 1304 | 1.36 | 254.695 | 0.85 | 0.016 | 257.866 | 0.665 | 0.012 | 0.505 | 0.483 | 0.368 | 0.144 |
| 1503 | 1.41 | 23.511 | 1.61 | 0.029 | 136.216 | 0.186 | 0.018 | 0.1 | 0.043 | 0.1 | 0.066 |
| 1606 | 1.46 | 175.342 | 0.62 | 0.036 | 351.572 | 0.526 | 0.023 | 0.1 | 0.048 | 0.126 | 0.048 |
| 1302 | 1.58 | 266.753 | 0.57 | 0.017 | 359.864 | 0.481 | 0.031 | 0.16 | 0.048 | 0.1 | 0.047 |
| 1011 | 1.61 | 730.53 | 0.67 | 0.012 | 719.768 | 0.729 | 0.014 | 0.111 | 0.056 | 0.437 | 0.338 |
| 1616 | 1.81 | 1154.46 | 1.33 | 0.097 | 1639.48 | 0.761 | 0.122 | 0.79 | 0.032 | 0.1 | 0.204 |
| 912 | 2.28 | 1917.44 | 0.72 | 0.022 | 7816.41 | 0.494 | 0.008 | 0.1 | 0.048 | 0.825 | 0.03 |

(\*) These subjects had either a STAGE I estimate of $\rho_i = 0$ or $\rho_i > 4$ . For those subjects we could not generate distributions of valuations for STAGE II that would adhere to our requirement to limit the number of trials with FOSD violations (when $\rho_i \rightarrow 0$ ), or without having to censor a very large tail of the Pareto distribution (when $\rho_i > 4$ ). Instead, for these subjects we plugged-in $\rho_i = 1$ to generate the distributions for STAGE II.
(\*\*) Subjects who had >20 FOSD violations in at least one of the treatments.

**Table S5.**
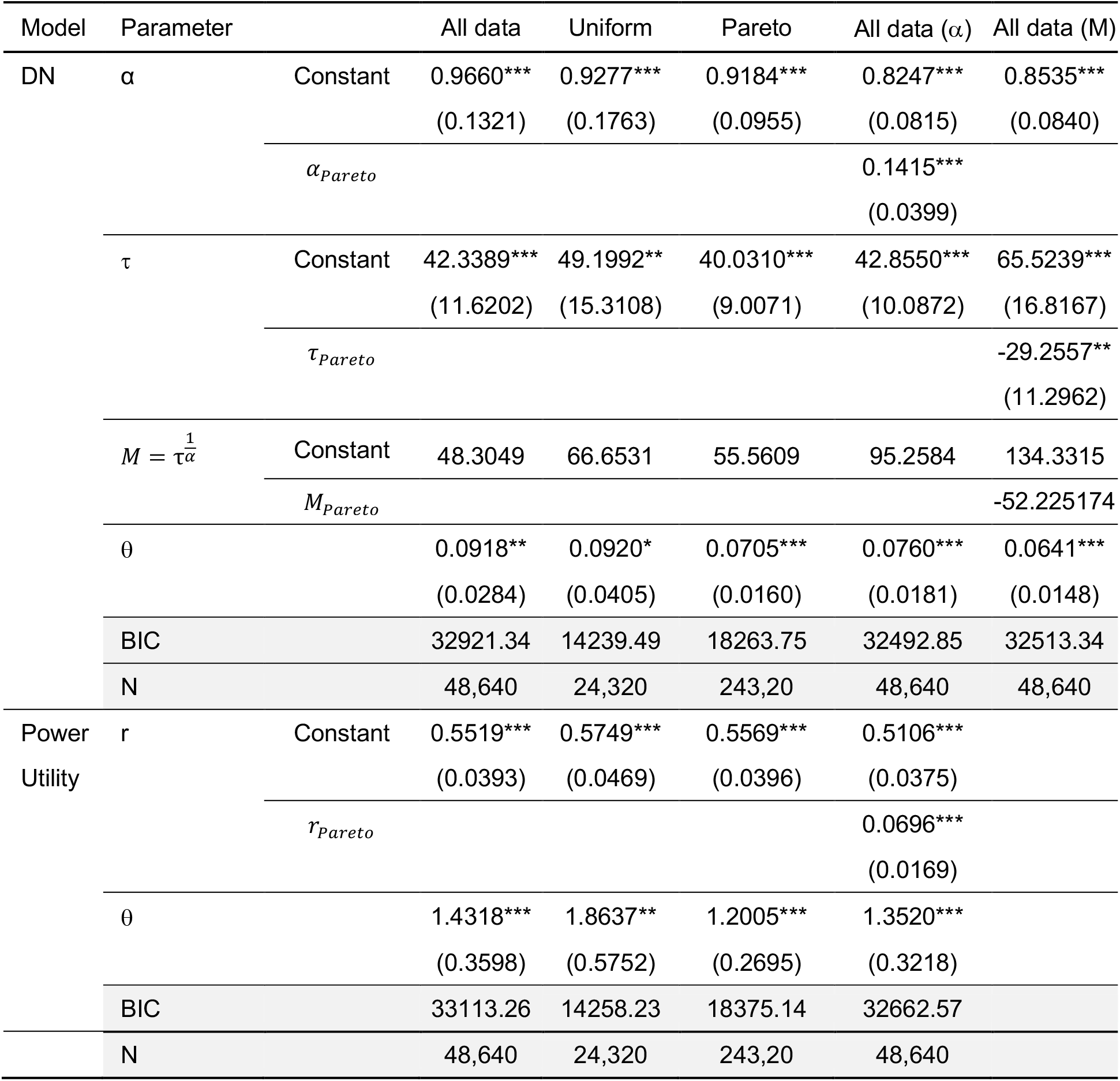
Pooled estimates, dollar space. In practice, to allow a better identification of the model parameters, we estimated the parameter *τ*, such that *τ* = *M*^*α*^. We recovered *M* post-hoc by simply plugging-in *τ* and *α* into the equation. Standard errors in parentheses, + p<0.1, * p<0.05, ** p<0.01, *** p<0.001.

## Footnotes

1 We note that the flexible power utility model nests within its parameterization a linear, a concave, and convex encoder.

2 The center (medians) of the distributions depended on subjects’ subjective valuation of dollar amounts (*ρ* parameter). In the Pareto distribution, the smallest median was $11.45 and the highest was $33.58. Likewise, in the uniform distribution, the smallest median was $22.85 and the highest was $41.53. Thus, determining a range of $9-42 included the distributions’ centers for all the subjects in our sample. See Table S3 for an alternative definition that included a smaller range.

3 Options in the uniform environment had on average higher value difference, thus responses in this environment were more accurate (Fig. S2) and less noisy (Fig. S3). Therefore, we only compare BIC scores of the two models within the same environment, and do not compare the models across the two environments.

4 This is because 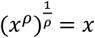.

5 Subjects also completed additional 640 trials with six-option choice sets with lottery valuations drawn either from a Pareto Type III or uniform distributions. Thus, in total, in each environment subjects encountered two 320 choice blocks. The six-option blocks were designed to examine another research question that is beyond the scope of the current study and will be reported in a separate paper. Blocks were presented in an order randomized across subjects but on a given day, all blocks were drawn from the same distribution. Payments for STAGE II included a realization of one choice from each of the two sessions, and could be drawn either from the two-options sets or from the six-options sets.

6 We set the location parameter *μ*_i_ = 0 to match the lower bound of the uniform distribution, and to avoid negative valuations.

## Notes

### Competing Interest Statement

The authors have declared no competing interest.

